# Evolution of mutation rates in rapidly adapting asexual populations

**DOI:** 10.1101/062760

**Authors:** Benjamin H. Good, Michael M. Desai

## Abstract

Mutator and antimutator alleles often arise and spread in both natural microbial populations and laboratory evolution experiments. The evolutionary dynamics of these mutation rate modifiers are determined by indirect selection on linked beneficial and deleterious mutations. These indirect selection pressures have been the focus of much earlier theoretical and empirical work, but we still have a limited analytical understanding of how the interplay between hitchhiking and deleterious load influences the fates of modifier alleles. Our understanding is particularly limited when clonal interference is common, which is the regime of primary interest in laboratory microbial evolution experiments. Here, we calculate the fixation probability of a mutator or antimutator allele in a rapidly adapting asexual population, and we show how this quantity depends on the population size, the beneficial and deleterious mutation rates, and the strength of a typical driver mutation. In the absence of deleterious mutations, we find that clonal interference enhances the fixation probability of mutators, even as they provide a diminishing benefit to the overall rate of adaptation. When deleterious mutations are included, natural selection pushes the population towards a stable mutation rate that can be suboptimal for the adaptation of the population as a whole. The approach to this stable mutation rate is not necessarily monotonic, and selection can favor mutator and antimutator alleles that overshoot the stable mutation rate by substantial amounts.

## INTRODUCTION

DNA replication occurs with extremely high fidelity, despite taking place in the noisy environment of the cell. For example, laboratory strains of *E. coli* produce a point mutation at a rate of roughly one nucleotide per ten billion copied (Lee et al., 2012; Wielgoss et al., 2011), which implies that hundreds of generations can elapse before a single mutation is introduced into the genome. To achieve such low error rates, bacteria and eukaryotes employ a complex array of cellular machinery, which must be maintained by natural selection.

One explanation for the low observed error rates is that the genome encodes a large number of functions, all of which are essential for survival. Low mutation rates could then emerge from hard selection against these lethal errors. But in practice, observed mutation rates lie far below the levels that would quickly result in extinction. This is evident from the fact that mutator strains, whose mutation rates are 10- to 1000-fold higher than the wild-type, can be propagated for thousands of generations in the laboratory without significant loss of viability (McDonald et al., 2012; Wiser et al., 2013). This suggests that mutation rates are not solely maintained by hard selection, but also by more direct evolutionary competition between strains with different mutation rates. These variants feel the effects of natural selection indirectly, by being linked to other mutations that directly influence fitness.

Strains with higher mutation rates are more likely to be linked to deleterious mutations, and will therefore experience an effective fitness cost. This cost can be measured in head-to-head competitions between mutator and wildtype strains (Chao and Cox, 1983; Chao et al., 1983; Gentile et al., 2011; Giraud et al., 2001; Thompson et al., 2006; Tröbner and Piechocki, 1981). In the absence of beneficial mutations, natural selection will therefore act to decrease the mutation rate, until it is eventually balanced by genetic drift, mutation, or other physiological costs. A large body of previous theoretical work has explored these dynamics (Dawson, 1998, 1999; Desai and Fisher, 2011; James and Jain, 2015; Johnson, 1999a; Kimura, 1967; Liberman and Feldman, 1986; Lynch, 2008, 2011; Soderberg and Berg, 2011).

When beneficial mutations are available, lineages with higher mutation rates are also more likely to produce and hitchhike with a successful beneficial variant. In the absence of deleterious mutations, natural selection will therefore act to increase the mutation rate, until the supply of beneficial mutations is eventually exhausted. In accordance with these expectations, mutator alleles are often found to spontaneously arise and spread in rapidly adapting microbial populations in the laboratory (Notley-McRobb et al., 2002; Pal et al., 2007; Shaver et al., 2002; Sniegowski et al., 1997; Voordeckers et al., 2015), and are often correlated with pathogenic lifestyles in the wild (Bjorkholm et al., 2001; del Campo et al., 2005; Denamur et al., 2002; Giraud et al., 2002; Labat et al., 2005; LeClerc et al., 1996; Matic et al., 1997; Oliver et al., 2000; Prunier et al., 2003; Richardson et al., 2002; Watson et al., 2004).

In a few cases, laboratory populations that have previously fixed a mutator allele have been shown to evolve a lower mutation rate later in the experiment, suggesting that mutation rate evolution is an ongoing process (McDonald et al., 2012; Notley-McRobb et al., 2002; Tröbner and Piechocki, 1984; Turrientes et al., 2013; Wielgoss et al., 2013). Because these populations often continue to increase in fitness during this process (Wielgoss et al., 2013), the success of mutator and antimutator alleles must depend on the interplay between beneficial and deleterious mutations, rather than the complete cessation of adaptation. However, our understanding of this interplay remains incomplete. A few simulation studies have shown how these effects can in principle favor either increases or decreases in mutation rates, depending on the specific population parameters (Taddei et al., 1997; Tenaillon et al., 2000, 1999; Travis and Travis, 2002). Some analytical progress has also been made in the case where beneficial mutations are rare, and occur one-by-one in discrete selective sweeps (Andre and Godelle, 2006; Desai and Fisher, 2011; Gillespie, 1981; Ishii et al., 1989; Johnson, 1999b; Leigh, 1970, 1973; Painter, 1975; Tanaka et al., 2003; Wylie et al., 2009) However, our understanding is much more limited in larger populations where beneficial mutations are more common, and multiple adaptive lineages must compete with each other for fixation. This is the regime where the indirect effects of linked selection are likely to be maximized, and it is also the most relevant for understanding the microbial evolution experiments where mutation rate modifiers have been observed to spread. In this clonal interference regime, even the most basic questions about this process remain unanswered: exactly how large can the deleterious load be before it effectively selects against a mutator allele? And how does this depend on the population size, the supply of beneficial mutations, and the magnitude of the mutator phenotype?

Here, we address these questions in the context of a simple model of adaptation, which is motivated by recent microbial evolution experiments. In particular, we focus on a regime in which deleterious mutations have a negligible impact on the rate of adaptation, even though they may have a large effect on the fates of mutation rate modifiers. Within this model, we calculate the fixation probability of an allele that changes the mutation rate by a factor *r* (with *r* > 1 corresponding to mutator alleles and *r* < 1 corresponding to antimutators). We consider both small changes in mutation rate (*r* ∼ 1) as well as substantial changes (where | log *r*| ≫ 1).

We find that clonal interference amplifies the beneficial advantage of mutator alleles, even as it constrains the overall rate of adaptation. For large *r* this can be a substantial effect, resulting in greater than 100-fold increases in the probability of fixation. However, clonal interference also amplifies the effects of the deleterious load, resulting in a much narrower window of mutator favorability. We show that this window depends most strongly on the strengths of typical driver mutations, and relatively weakly on the rate at which beneficial mutations arise. With the steady production of such modifier alleles, the mutation rate will evolve towards a stable point at which neither mutator or antimutator alleles are favored. Surprisingly, we find that the approach to this stable point is not necessarily monotonic, even in the absence of epistasis. Instead, alleles that overshoot the stable point can be positively selected, and selection must then act to change the mutation rates in the opposite direction.

## MODEL

Our goal is to understand how beneficial and deleterious mutations influence the fates of mutation rate modifiers in rapidly adapting populations. We focus on the simplest possible model in which we can address this question. Specifically, we consider an asexual haploid population of constant size *N* (our analysis also applies to diploids when all mutations have intermediate dominance effects). We assume beneficial and deleterious mutations occur at genome-wide rates *U_b_* and *U_d_* respectively. We define *U* = *U_b_* +*U_d_* as the total mutation rate and 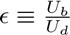as the ratio of beneficial to deleterious mutation rates. We focus on adapting populations, and consider both the successional-mutation regime (where beneficial mutations are rare) and the clonal interference regime.

For most of our analysis we assume that beneficial mutations all confer the same fitness advantage *s_b_*, though we also comment on how our results can be extended to the case where beneficial mutations have a distribution of fitness effects. We assume throughout that there is no macroscopic epistasis among beneficial mutations, so that *U_b_* and *s_b_* remain fixed as the population adapts. We make no specific assumptions about the fitness effects of deleterious mutations, other than that they are *purgeable* — that is, deleterious mutations are sufficiently costly that they are unlikely to fix and do not significantly reduce the overall rate of adaptation of the population. We describe the technical conditions required for this to be true and the qualitative implications of non-purgeable deleterious mutations in more detail in the Discussion.

Our goal is to analyze the fate of a modifier allele that changes the overall mutation rate *U* by a factor *r* (while leaving ∈ and *s_b_* unchanged). For simplicity, we assume that this allele has no direct influence on fitness, although in the Discussion we show how this assumption can be relaxed. These modifier alleles are produced at rate *μ* from the wildtype background, and we assume that *μ* is sufficiently small that the modifiers can be treated independently. In this case, the substitution rate of the modifier is 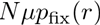, where *p*_fix_(*r*) is the fixation probability of a single modifier individual. A natural measure of how selection “favors” or “disfavors” the modifier can be obtained from the scaled fixation probability, *N*_*p*_fix__(*r*). When *N*_*p*fix_ > 1, the allele is favored by selection, since it substitutes more rapidly than a neutral allele; conversely, the allele is disfavored when *N*_*p*_fix__ < 1. This scaled fixation probability will be our primary focus in the analysis below. In particular, we are interested in determining the parameters where *N*_*p*_fix__ transitions between favored (*N*_*p*_fix__ > 1) and unfavored (*N*_*p*_fix__ < 1).

We are primarily interested in situations applicable to microbial populations, and particularly to laboratory evolution experiments. This motivates certain technical assumptions we describe in the Analysis below. It also motivates our focus on asexual populations where recombination can be neglected. This asexual case is particularly relevant because when recombination is absent, modifiers of mutation rate remain perfectly linked to beneficial or deleterious mutations they generate, and hence experience the strongest indirect selection pressures. In principle, we could also apply our analysis to physically linked regions within sexual genomes (“linkage blocks”) that are unlikely to recombine on the appropriate timescales (Good et al., 2014; Neher et al., 2013; Weissman and Hallatschek, 2014). However, the scale of these linkage blocks is a complex problem, and understanding these effects is beyond the scope of this study.

In the remaining sections, we calculate *N*_*p*_fix__(*r*) as a function of the underlying model parameters, and then we use these predictions to analyze the dynamics of mutation rate evolution.

## THE SUCCESSIONAL MUTATIONS REGIME

To gain intuition into the forces that determine *N*_*p*_fix__(*r*), it is useful to start by considering the simplest case, where deleterious mutations can be neglected and adaptation proceeds by a sequence of discrete selective sweeps. This “successional mutations” regime applies whenever beneficial mutations are sufficiently rare that they seldom segregate within the population at the same time. This will be true provided that 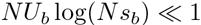 (Desai and Fisher, 2007). In this regime, beneficial mutations are produced at rate *NU_b_*, and establish with probability 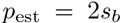 Selective sweeps then occur as a Poisson process with rate

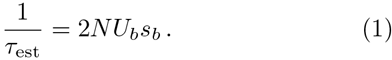

When a successful beneficial mutation establishes, it starts to grow deterministically and requires roughly 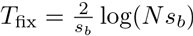 generations to complete its sweep. By definition, this time is small compared to the time between sweeps, so the rate of adaptation is just *v* = *s_b_*/*τ*_est_. A modifier allele would increase or decrease this rate by factor of *r*, if it was able to fix. We now analyze the dynamics of this process.

When a modifier allele first arises, it starts at an initial frequency *f_m_*(0) = 1/*N*, and it will then drift neutrally until the next selective sweep occurs. Most of the time, the lineage will fluctuate to extinction within the first few generations. However, with probability ~ 1/*t*, it will survive for ~ *t* generations and reach size *f_m_*(*t*) ~ *t*/*N* (Fisher, 2007). In the absence of selective sweeps, the mutation modifier lineage could only fix by drifting to fixation; this happens with probability ~ 1/*N* and occurs over a timescale of order ~ *N* generations. Thus, depending on how this drift timescale *N* compares with the sweep timescales *τ*_est_ and *τ*_est_/*r*, two distinct types of fixation dynamics can emerge.

When *τ*_est_ and *τ*_est_/*r* are both much much larger than *N*, then the fate of the modifier lineage will be determined long before the next selective sweep occurs. Drift is the dominant evolutionary force, and *p*_fix_(*r*) = 1/*N*.

In contrast, if either *τ*_est_ and *τ*_est_/*r* are small compared to *N*, the fixation process is instead controlled by genetic draft. In this case, the modifier lineage will only fix if it is lucky enough to produce and hitchhike with the next selective sweep. Since we are primarily interested in understanding these effects, we will focus on this regime for the remainder of our analysis.

The next selective sweep is produced by two competing Poisson processes, corresponding tothe beneficial mutants from the wildtype and modifier lineages. These respectively produce sweeps at rates

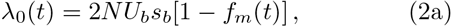

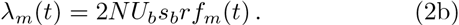

The probability that the next sweep is produced by the modifier lineage is then simply

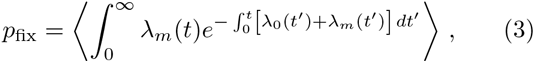

where the angled brackets denote an average is over the random lineage trajectory *f_m_*(*t*). In the regime we are considering, the modifier lineage will remain rare until the next selective sweep occurs. The trajectory *f_m_*(*t*) is therefore described by the Langevin dynamics,

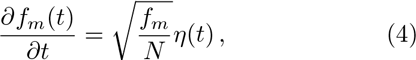

where *η*(*t*) is a Brownian noise term (Gardiner, 1985). The solution of Eq. (3) can be complicated, since both the overall rate of sweeps and their probability of arising in the modifier lineage depend on the random trajectory *f_m_*(t). We present an exact solution of these equations in Appendix A, using techniques developed by Weissman et al. (2009).

For the present purposes, however, it will be more useful to focus on an approximation. If the modifier lineage stays sufficiently rare that it does not influence the overall rate of sweeps,then we can approximate 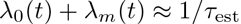. Equation (3) then reduces to

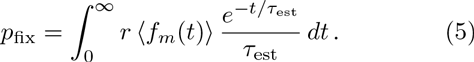

In other words, the fixation probability is given by the average size of the lineage at the time of the next sweep, and this time is unaffected by the presence of the modifier. In this case, the modifier lineage is neutral and 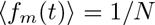, so that

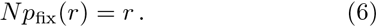

Thus, we see that the modifier fixation probability (like the proportional change in the rate of adaptation) is independent of the population size, the mutation rate, and the fitness benefits of the driver mutations.

To derive Eq. (6), we made the approximation that 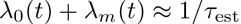. Substituting our expressions for λ_0_ and λ_m_(*t*) in Eq. (2), we see that this is equivalent to the assumption that (*r* — 1)*f_m_*(*t*) ≪ 1. However, since *f_m_* and *t* are both random variables, we cannot determine the validity of this assumption based on the average values alone. We must instead consider the *typical* dynamics that contribute to the average in Eq. (5): with probability ~ 1/*τ*_est_, the modifier lineage survives for *τ*_est_ generations and reaches size 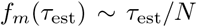. This ensures that the average size of the lineage is just 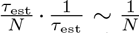, as expected. It also shows that the typical values of (*r* — 1)*f_m_*(*t*) will remain small compared to one provided that *rτ*_est_ ≪ N. For simplicity, we will assume that this condition holds for the remainder of our analysis. For moderate *r*, this is actually the same assumption we already made in focusing on the genetic-draft regime. For large *r* it imposes an upper limit *r* | *N*/*τ*_est_ ≫ 1 on the range of modifier effects that we consider. However, this is not a fundamental problem for the analysis; we consider such large-effect modifiers in Appendix A.

### Incorporating purgeable deleterious mutations

A similar picture applies in the presence of strongly deleterious mutations. In particular, we assume that the typical fitness costs are larger than *s_b_*, so that a driver mutation can only fix in a background that is free of deleterious mutations. If the costs are also much larger than *U_d_*, then the vast majority of the population will be of this mutation-free type. We refer to these as *purgeable* mutations, and we will assume that all deleterious mutations are purgeable in the analysis that follows.

When a new beneficial mutation occurs, the wildtype will be in mutation-selection balance with respect to its deleterious mutations, so the mean fitness of the population is —*U_d_*. Even if it arises on a mutation-free background, the new beneficial lineage will continue to produce loaded individuals at rate *U_d_*. This exactly cancels the growth advantage of the mean fitness of the population, so that the establishment probability is still just 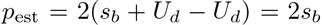. Since most genetic backgrounds are in this unloaded state, the overall rate of sweeps is unchanged from before. This means that if a modifier allele were to fix, it would still change the rate of adaptation by a simple factor of *r*.

However, while the deleterious mutations do not affect the rate of adaptation, they can still have a large effect on the modifier lineage while it is rare. The modifier produces doomed lineages at rate rU_d_, which is not completely cancelled by the mean fitness of the wild-type. Instead, mutator alleles will feel an effective cost of magnitude *U_d_*(*r* – 1), while antimutators will experience an effective fitness advantage. Generalizing Eq. (4), the mutation-free portion of the modifier lineage will then evolve as

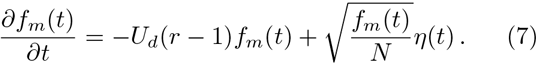

Beneficial mutations produced by this lineage will likewise have an effective fitness *s_b_* – *U_d_*(*r* – 1), and will establish with probability

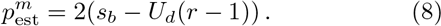

In exactly the same manner as before, the next selective sweep is produced by two competing Poisson processes with rates

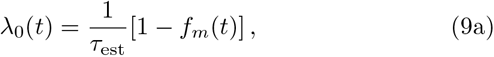

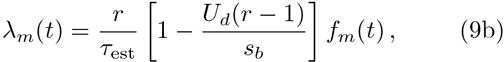

and the fixation probability is given by the average in Eq. (3). If we again make the assumption that the modifier lineage has a negligible influence on the overall sweep rate 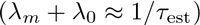, this reduces to

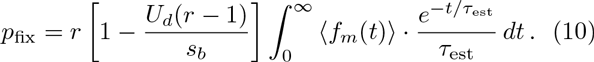

In this case, the average lineage size is now 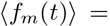
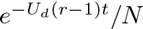 and we obtain

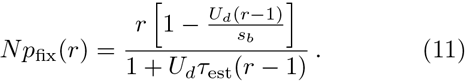

The validity of this expression depends on our assumption that 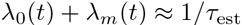. In the same manner as above, we can establish the region of validity of this approximation by considering the typical dynamics behind the averages in Eq. (10). These will depend on the sign of *U_d_*(*r* − 1).

For a mutator allele (*r* > 1), the modifier lineage will feel an effective fitness cost *U_d_*(*r* − 1). If this cost is small compared to 1/*τ*_est_, then selection will barely have time to influence *f_m_*(*t*) before the next sweep occurs, and mutators will continue to fix with probability *N*_*p*_fix__ ~ *r*. On the other hand, if *U_d_*(*r* − 1) ≫ *τ*_est_, then this fitness cost exponentially suppresses the probability that the mutator survives until the next sweep. In particular, the mutator will survive for *t* generations with probability 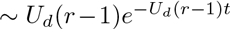, and provided that it does so, it will no longer grow indefinitely, but will instead reach a maximum size of order 1/*NU*_d_(*r* − 1). If the time between sweeps was deterministic, this would simply suppress the fixation probability by a factor 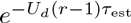. However, this is not actually the case, since the next sweep is generated by the random Poisson process above. Thus, the intervals between sweeps are not only random, but they are drawn from an exponential distribution, which has important consequences for the dynamics. The broad distribution of *t* does not lead to a simple exponential suppression of *N*_*p*fix_(*r*), but rather a power law decay 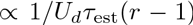, which is dominated by anomalously early sweeps for which 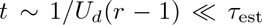. For a strong-effect mutator (*r* ≫ 1), this power-law decay exactly balances the lineage’s increased beneficial mutation rate, so that the *r*-dependence of *p*_fix_ is primarily mediated through the reduced establishment probability, 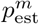.

In contrast, antimutators will have an effective fitness advantage relative to the wildtype, of magnitude 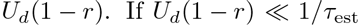, this fitness advantage will still have a negligible influence on *N*_*p*fix_(*r*). However, when *U_d_*(1 − *r*)*τ*_est_ ≫ 1, the antimutator will establish with probability *U_d_*(1 − *r*) and will start to grow deter-ministically as 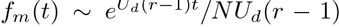. Again, if the time to the next sweep was deterministic, this would result in a simple exponential enhancement of *N*_*p*fix_. But because *t* is exponentially distributed, it interacts with the exponential growth of the lineage in a strong way. In particular, as *U_d_*(1 − *r*)*τ*_est_ approaches 1, the average in Eq. (11) will be dominated by anomalously late sweep times for which *f_m_*(*t*) is no longer rare, and our approximation that λ_0_ + λ_m_ ≈ 1/*τ*_est_ breaks down. Thus, Eq. (11) will only be valid provided that 1 − *U_d_*(1 − *r*)*τ*_est_ ≫ 1/log(*N*/*τ*_est_). Outside this regime, the antimutator lineage will appear to sweep to fixation on its own, withoutthe help of an additional driver mutation.

**FIG. 1.**
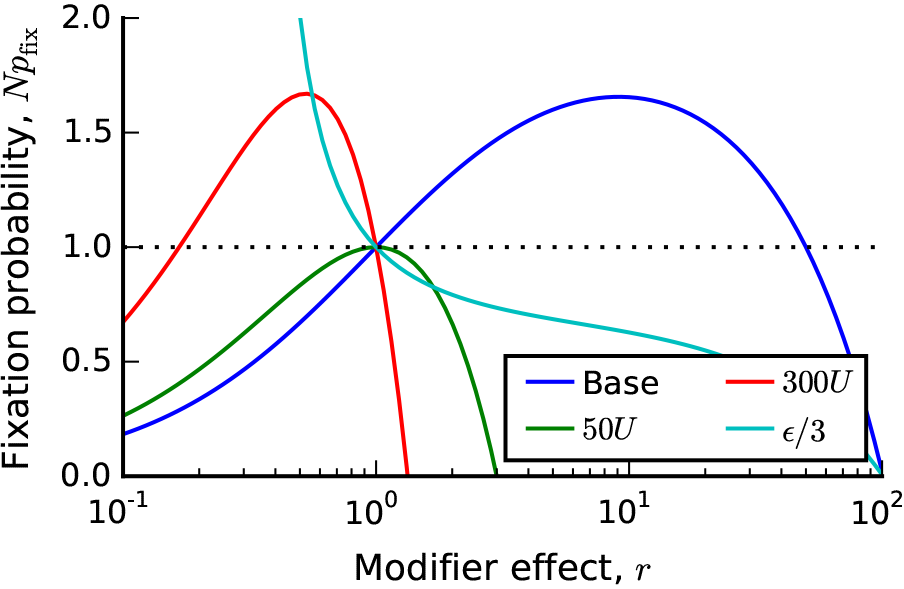
The fixation probability of a mutation rate modifier in the successive mutations regime. Solid lines depict Eq. (11) for four sets of parameters, which illustrate the four characteristic shapes of *N*_*p*fix_(*r*). In all four cases, the base parameters are *N* = 10^7^, *s_b_* = 10^-2^, *U* = 10^-4^ and ∈ = 10^-5^, with modifications listed in the legend.

The fixation probability in Eq. (11) is illustrated in Fig. 1 for several choices of parameters. Its overall shape is determined by the key parameter

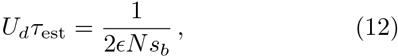

which depends only on *N*, *s_b_*, and ∈ ≡ *U_b_*/*U_d_*, and is independent of the overall mutation rate *U*. Depending on the value of U_d_T_est_, Eq. (11) takes on one of two characteristic shapes. If *U_d_τ*_est_ > 1, the fixation probability is a monotonically decreasing function of *r* (see Fig. 1), and mutators will never be favored to invade. Instead, antimutators will be positively selected. Note that this happens even though the antimutator actually causes a *decrease* in the overall rate of adaptation.

On the other hand, for *U_d_τ*_est_ < 1, the fixation probability takes on a paraboloid shape (Fig. 1). It crosses *N*_*p*flx_(*r*) = 1 at exactly two points: once at r = 1 and once at

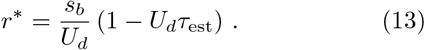

When *r\** > 1, modifiers will be favored in the range 1 < *r* < *r\**, and disfavored elsewhere (i.e., some mutators will be favored). Conversely, when *r\** < 1, modifiers will be favored in the range *r\** < *r* < 1 (i.e., some antimutators will be favored). It is also useful to rewrite *r\** in terms of the mutation rate *U_m_* = *Ur* of the modifier lineage:

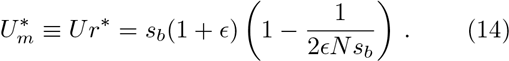

Note that this is always positive (since 1/2*N s_b_* ∈ < 1), and it is independent of *U*.

### Dynamics of mutation rate evolution

We are now in a position to analyze how mutation rates evolve over time. Imagine that we start with some particular values of *N*, *U*, ∈, and *s_b_*. This will correspond to a particular value of *τ*_est_ and 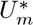. If *U*_*d*_*τ*_est_ > 1, mutators are disfavored and antimutator alleles will be positively selected instead. After an antimutator allele fixes, both *U_b_* and *U_d_* are reduced, but *U_d_τ*_est_ is unchanged, so natural selection will continue to select for lower mutation rates until this force is balanced by drift, mutational pressure, or other physiological costs.

On the other hand, if *U_d_τ*_est_ < 1 and 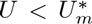, mutators will be favored provided that their resulting mutation rate is less than 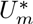. If a mutator allele fixes, *U_b_* and *U_d_* will be increased, but *U_d_τ*_est_ and 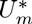 will remain constant. Thus, natural selection will continue to favor increased mutation rates until *U* reaches a special value,

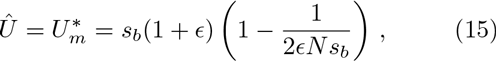

which is stable against further changes in the mutation rate. If instead the initial mutation rate *U* > *Û*, antimutators will be favored provided that their resulting mutation rate is greater than *Û* and natural selection will continue to favor increased mutation rates until *U* reaches *Û*. We describe these dynamics in more detail in the Discussion.

Of course, this analysis crucially depends on the assumption that any mutation rate increases will still lie within the successive mutations regime. This will be true provided that 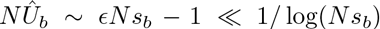. But in order for a nonzero stable mutation rate to exist in the first place, we previously required that 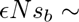 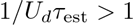. These two conditions can only be satisfied if 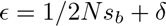, where

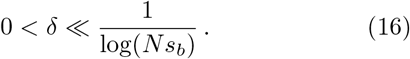

This is an extremely small region of parameter space, and it grows increasingly narrow as *N S*_*b*_ increases. As a result, the stable mutation rate in Eq. (15) is actually a rapidly varying function of *N*, *S*_*b*_, and ∈. For larger values of ∈ that violate the stringent constraints in Eq. (16), mutator alleles will generically drive mutation rates into regimes where selective sweeps begin to interfere with each other. We now turn to an analysis of this case.

## THE CLONAL INTERFERENCE REGIME

When multiple beneficial mutations segregate at the same time, many potential drivers are lost due to competition with other, fitter genetic backgrounds. This reduces the rate of *successful* drivers in a way that depends on the relative values of *N s_b_* and *NU_b_*. In the regime most relevant for our current study, Desai and Fisher (2007) have shown that *τ*est is reduced to

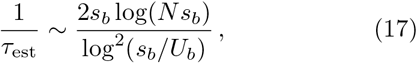

which is valid provided that 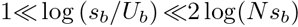 and 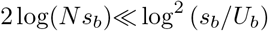. Similar expressions canbe obtained for other parameter regimes, all of which share the weak dependence on *NU_b_* (Fisher, 2013). Since adaptation is no longer mutation-limited, one might guess that mutators will be less strongly favored in this regime. However, previous simulation studies (Tenaillon et al., 1999) and heuristic reasoning (Desai and Fisher, 2011) suggest that the opposite can actually be true: clonal interference enhances the fixation of mutator alleles, even as they provide a diminishing overall benefit for the rate of adaptation.

In the following subsection, we introduce a traveling-wave formalism for calculating *N*_*p*fix_(*r*) in the presence of clonal interference. Before doing so, however, it will be useful to consider this process from a heuristic perspective. This will allow us to identify the key forces and parameters involved, and will provide intuition for the more rigorous analysis that follows.

### Heuristic analysis

In the absence of deleterious mutations, clonal interference alters the dynamics of fixation in two main ways. First, successful mutations can only occur in the most highly-fit genetic backgrounds in the population. These individuals lie in the extreme right tail or “nose” of the population fitness distribution, which steadily increases in fitness as the population adapts. In the regime described by Eq. (17), the relative fitness of the nose is given by

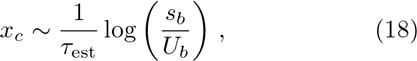

which is much larger than the size of a single driver mutation. These individuals have already acquired *q* = *x_c_*/*s_b_* ≫ 1 more adaptive mutations than the average individual, but they are not yet destined to fix. The reason is that there are still enough individuals in the nose that they will collectively produce *multiple* additional driver mutations in the next *τ*_est_ generations, and these will occur in a relatively narrow time window Δ*t* ≪ T_est_ (Desai and Fisher, 2007). By definition, all but one of these mutations must eventually be outcompeted. But this means that the process of fixation within the nose takes place over *multiple* establishment intervals, each of length *τ*_est_.

Since *x_c_τ*_est_ ≫ 1, genetic drift is not directly relevant for most of this fixation process. Random fluctuations in the lineage sizes are still important, but these are now driven by *genetic draft*, which arises from slight differences in the relative order of the next round of driver mutations. During most of this process, the lineages founded by different driver mutations make up less than ~ 1/*q* of the total size of the nose. However, approximately once every ~ *q* establishments, an anomalously early driver mutation will occur and reach an 𝒪(1) fraction of the nose (Desai et al., 2013). This roughly coincides with a fixation event. In order to fix, a lineage must therefore (i) arise in the nose, (ii) persist for ~ *q* additional establishment intervals, and (iii) be lucky enough to hitchhike with the special “jackpot” driver event.

Mutation-rate modifiers can be incorporated into this picture in a straightforward way. The key quantities *τ*_est_ and *q* are only weakly dependent on the mutation rate. Provided that | log(*r*)*|* ≪ log(*s_b_*/*U_b_*), the modifier versions of these parameters will be similar to the wildtype, and the competition between these lineages will play out according to the same dynamics as above. The modifier is no more or less likely to arise in the nose compared to a neutral mutation. However, provided that it does occur in one of these special genetic backgrounds, a mutator lineage is *r* times more likely than a neutral lineage to generate an additional driver, while an antimutator lineage is *r* times less likely to do so. Since the modifier lineage must generate ~ *q* additional drivers to fix,its overall fixation probability is given by

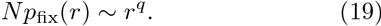

Thus, we see that clonal interference increases the fixation probability of mutators by ~ *q* factors of *r*. This increase can be substantial for large *r*, even when *q* ≈ 2 (see Fig. 2). From our discussion above, we see that this increase is primarily driven by the fact that *multiple* additional drivers are required for fixation.

In the presence of deleterious mutations, a mutator lineage again feels an effective cost of *U_d_*(*r* − 1), so it will tend to decline in frequency relative to the other individuals in the nose. In particular, when the next burst of driver mutations arises, the mutator lineage will have decreased in frequency by a factor of 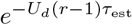, which makes it that much less likely to survive to the next round. Similarly, an antimutator lineage will have increased in frequency by an analogous factor of 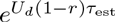. The overall fixation probabilitythen be-comes

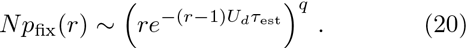

**FIG. 2.**
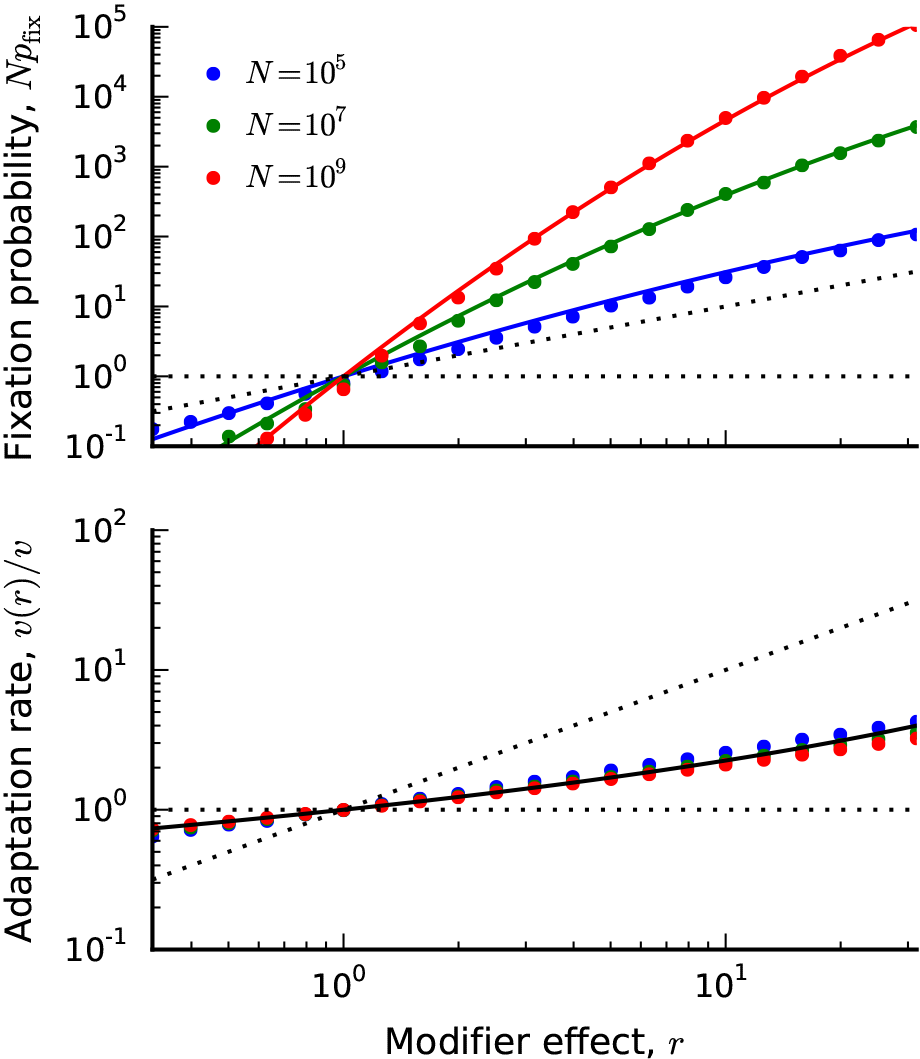
(Top) The fixation probability of a mutation rate modifier in the clonal interference regime when *U_d_* = 0. Symbols denote the results of forward-time solutions (described in Appendix C) for *s_b_* = 10^-2^, *U_b_* = 10^-5^, and *N* ∈ {10^5^,10^7^, 10^9^}. Solid lines denote the theoretical predictions in Eq. (38). For comparison, the successive mutations prediction (*N*_*p*fix_ ≈ *r*) and neutrality (*N*_*p*fix_ ≈ 1) are shown in the dashed lines. (Bottom) The rate of adaptation of a successful modifier lineage, relative to that of the wildtype, for the same set of populations above. The solid black line denotes the asymptotic prediction, 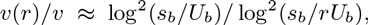, which is independent of N. For comparison, the successive mutations prediction (*v*(*r*)/*v* ≈ *r*) and no change (*v*(*r*)/*v* ≈ 1) are shown in the dashed lines.

We discuss the regimes of validity of this expression in our more rigorous analysis below. For the purposes of our heuristic analysis, it will be more useful to focus on the implications of Eq. (20).

Similar to the selective sweeps case, the direction of selection in Eq. (20) is again determined by the product *U_d_τ*_est_. There are three characteristic regimes of behavior. When *U_d_τ*_est_ ≪ 1, mutators will be favored provided that *U_m_* = *Ur* is less than a maximum value,

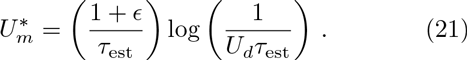

Conversely, when *U_d_τ*_est_ ≫ 1, antimutators will be fa-vored above a minimum value

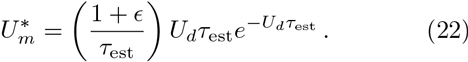

Similar behavior is obtained for *U_d_τ*_est_ is close to one, except that 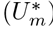 is now given by

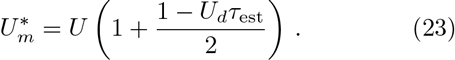

When *U_d_τ*_est_ = 1, the range of favorable modifiers vanishes, and mutators and antimutators are both selected against. The evolutionarily stable mutation rate is therefore given by

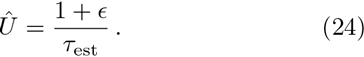

Note that this is actually an implicit relation for *Û*, since *τ*_est_ depends on *Û_b_* = ∈Û/(1 + ∈). Substituting our expression for *τ*_est_ in Eq. (17) and solving for *Û*, we find that

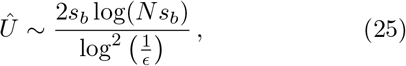

where the region of validity for Eq. (17) [and hence Eq. (25)] now becomes

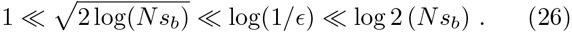

This is still a restrictive parameter range for e, although it is much broader than Eq. (16) in the successive mutations regime, and it grows larger with increasing *N s_b_*.

Provided that these conditions are met, we see that the stable mutation rate in Eq. (25) is only weakly dependent on *N* and ∈, and is much more strongly influenced by *s_b_*. This contrasts with the behavior we found in the successive mutations regime. In addition, we see that unlike in the successive mutations regime, the stable mutation rate (*Û*) does not necessarily coincide with the maximally permitted mutator or antimutator allele 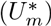 when *U* ≠ *Û*. This can have important consequences for the dynamics of mutation rate evolution in this regime. If the mutation rate starts far above or below *Û*, natural selection can favor modifier alleles that overshoot the stable value by a substantial amount, which can lead to a nonmonotinic approach to the stable point. We will revisit these dynamics in more detail in the Discussion.

### Formal analysis

We now turn to a more formal derivation of the results described above. To do so, we make use of “traveling-wave” formalism developed in previous work (Fisher, 2013; Good and Desai, 2014; Good et al., 2012; Hal-latschek, 2011; Neher and Shraiman, 2011; Neher et al., 2010). This formalism focuses on the distribution of relative fitness within the population, which we denote by *f*(*x*), and the fixation probability of a (wildtype) individual with relative fitness *x*, which we denote by *w*(*x*). In the absence of mutator or antimutator alleles, we and others have previously shown that *f*(*x*) and *w*(*x*) satisfy the partial differential equations,

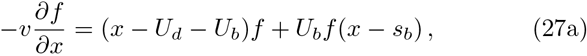

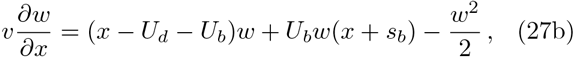

where *v* ≡ *s_b_*/*τ*_est_ is the average rate of adaptation (Good and Desai, 2014). Note that Eq. (27) implicitly assumes that the deleterious mutations are purgeable, i.e., they do not fix and consitute a negligible fraction of the population. The rate of adaptation (or equivalently, *τ*_est_ ≡ *s_b_*/*v*) is set by the self-consistency condition,

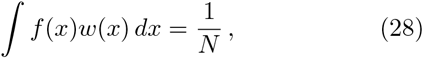

which is just another way of saying that the fixation probability of a neutral mutation must be equal to 1/*N*. For a detailed derivation of these equations, see Good and Desai (2014). Note that because we have assumed that deleterious mutations are purgeable, *U_d_* enters into Eqs. (27) and (28) only as an overall shift in the mean fitness of the population. If we measure fitnesses using the shifted variable 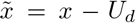 (i.e., fitness relative to the average *unloaded* individual), then the evolution equations revert back to the purely beneficial case. We will adopt this convention from now on.

Even in the simplified model of Eq. (27), there are many possible parameter regimes that one may consider (Fisher, 2013). Here we will focus on a particular approximate solution which is thought to be relevant for many microbial evolution experiments. In this regime, the fitnesses of unloaded individuals are approximately normally distributed,

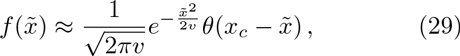

with a sharp cutoff at 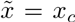. This corresponds to the “nose” of the fitness distribution described above. Meanwhile, the survival probability 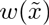 can be approximated by the piecewise form

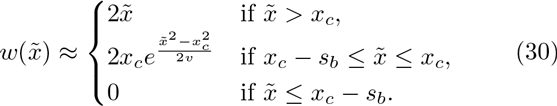

In this context, *x_c_* can also be thought of as an *interference threshold*. Lineages that are at relative fitness *x* > *x_c_* will fix provided they survive drift and establish; this occurs with probability 2*x*. Below *x_c_* the fixation probability drops off rapidly because lineages can establish but still be lost to interference. Once a lineage is more than *s_b_* below *x_c_*, it can essentially never catch up with the remaining lineages at the nose, and it will therefore have a negligible fixation probability. The location of *x_c_* is determined by the auxiliary condition

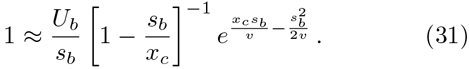

After substituting our expressions for *f*(*x*) and *w*(*x*) into Eq. (28), the self-consistency condition becomes

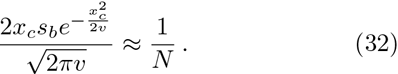

Together with Eq. (31), this completely determines *v* and *x_c_* as a function of the underlying parameters. Asymptotic formulae for these quantities are given in Eqs. (17) and (18); more accurate estimates can be obtained by solving Eqs. (31) and (32) numerically.

We have previously shown that this solution applies whenever 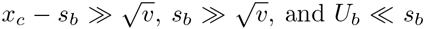. (Good and Desai, 2014). The first of these conditions says that the parents of successful lineages must be highly-fit (i.e., that clonal interference is indeed present). The second condition (which also implies the third) says that the vast majority of individuals in the population have roughly the same number of driver mutations (i.e., mutation pressure is not too strong). In terms of the underlying parameters, these conditions become 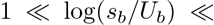 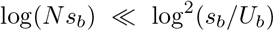, as we stated after Eq. (17) above.

Once we have set up this travelling-wave formalism, mutation-rate modifiers can be added in a straightforward way. Since the fate of any allele is determined while it is rare, we can neglect the effects of the modifier on the wildtype population, so that *f*(*x*) and *v* are unchanged from above. When a modifier allele occurs, it will arise on a genetic background whose fitness is drawn from *f*(*x*). We then introduce a new function *w_m_*(*x*) describing the fixation probability of the modifier allele as a function of its initial fitness. This function satisfies a similar equation as *w*(*x*), except with a different mutation rate:

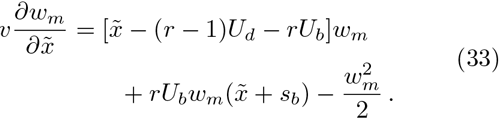

Thus, in addition to the increase in *U_b_*, we see that a mutator lineage experiences an effective fitness cost *U_d_*(*r*−1) corresponding to its increased deleterious load. This cost enters as a shift in the overall location of 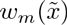, which we can account for by defining a shifted function, 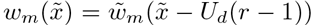. After this transformation, Eq. (33) is of the same form as Eq. (27b). Then, provided that the modifier mutation rate rU_b_ is still in the same regime as U_b_, the solution for the shifted function 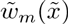 will have the same form as 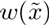,

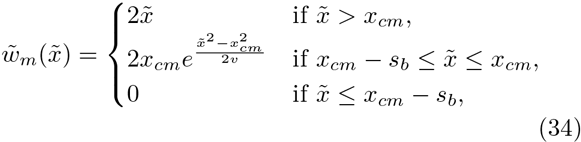

except with a different interference threshold *x_cm_* satisfies

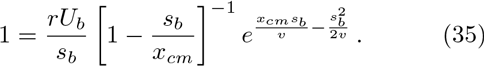

Like the wildtype interference threshold *x_c_*, the location of *x_cm_* is independent of *U_d_*. Since mutators are favored when *U_d_* = 0, we expect that *x_cm_* < *x_c_* whenever *r* > 1.

In other words, we expect that mutators have a lower interference threshold, since these lineages generate beneficial mutations more rapidly and, once in the nose, are therefore less likely to be lost to clonal interference.

In order for this solution to apply, it must satisfy the same conditions as the wildtype population: 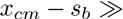 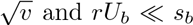.This will be true provided that x_cm_ is still close to x_c_, or equivalently, that the fractional difference *δ*_x_ = *x_cm_*/*x_c_* − 1 is small compared to 1. Given this assumption, we can divide Eq. (35) by Eq. (31) and solve for *δ_x_*:

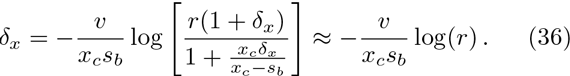

Since *x_c_s_b_*/*v* ~ log(*s_b_*/*U_b_*) in this regime, we see that this approximation will hold provided that | log(*r*)| ≪ log(*s_b_*/*U_b_*). This places an upper limit on the range of modifier effects that we can consider. But since *s_b_*/*U_b_* ≫ 1, this includes many realistically-large modifiers of order *r* ~ 100.

Given our solution for 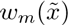, we can compute the marginal fixation probability of the modifier by averaging over all the fitness backgrounds that the modifier could have arisen on:

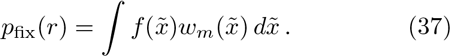

**FIG. 3.**
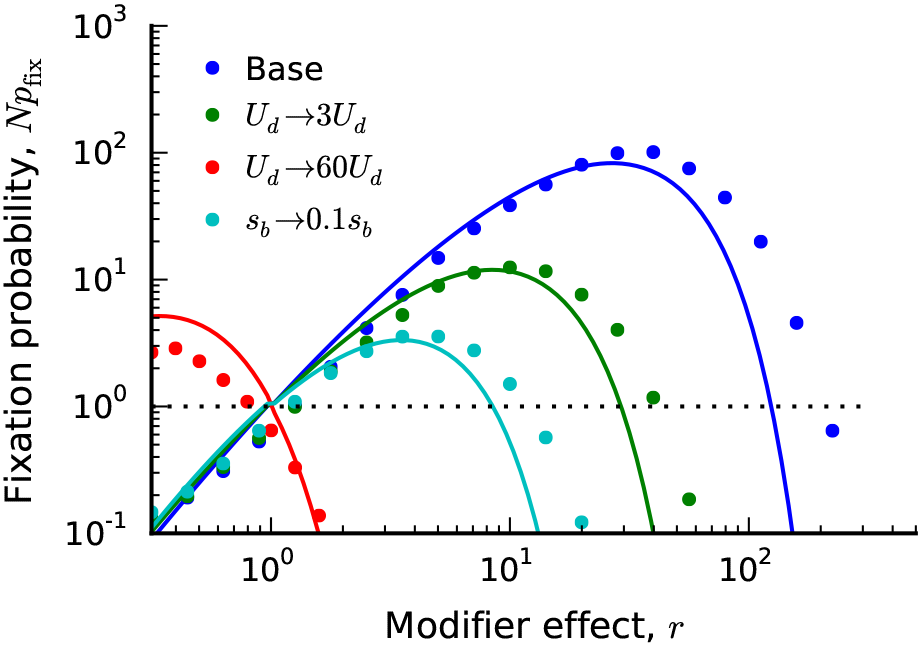
The fixation probability of a mutation rate modifier in the clonal interference regime when *U_d_* >0. Symbols denote the results of forward-time solutions with the base parameters *N* = 10^7^, *s_b_* = 10^-2^, *U_b_* = 10^-5^, *U_d_* = 10^-4^, and Sd = 10^-1^, with modifications listed in the legend. Solid lines denote the theoretical predictions in Eq. (38).

We evaluate this integral in Appendix B. We find that

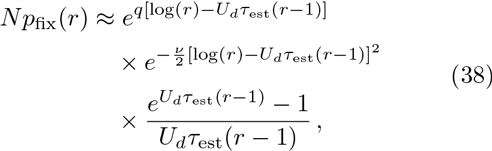

where we have employed the short-hand notation *q* = *x_c_*/*s_b_*, *τ*_est_ = *S_b_*/*v*, and *v* = 1/*s_b_τ*_est_. In the asymptotic regime we are considering, 1/q and v are both small parameters. To leading order, *Eq. (38)* converges to the simpler expression from our heuristic analysis, which we now recognize as the technical statement that

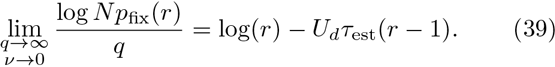

However, since the errors are multiplied by a large number *q* and exponentiated, the full version in Eq. (38) is often required for quantitative accuracy.

We illustrate the full expression in Eq. (38) and compare it to forward-time simulations in Figs 2 and 3. The accuracy is generally good, although there are some systematic deviations for large *r* when the effects of the deleterious load are particularly costly. These are ultimately the result of two factors: (i) inaccuracies in our approximate solution for *w*(*x*) for *x_c_* − *s_b_* < *x* < *x_c_* and (ii) the fact that log(*r*) is starting to approach log(*s_b_*/*U_b_*). More accurate approximations for *w*(*x*) derived by Fisher (2013) could be used to improve the quantitative accuracy for these parameters; we leave this for future work.

## DISCUSSION

In rapidly adapting populations, the fates of mutator and antimutator alleles depend on a careful balance between the benefits of hitchhiking and the cost of a deleterious load. Our analysis shows how the fixation probabilities of these mutation-rate modifiers depend on the population size, the beneficial and deleterious mutation rates, and the strength of selection. By searching for parameters where mutators and antimutators are both disfavored by selection, we can identify particular mutation rates *Û* which are stable under future evolution. At these mutation rates, the benefits of hitchhiking are exactly balanced by the costs of the deleterious load, so that neither mutators nor antimutators are favored. We now consider how a population will approach the stable mutation rates, and the implications of this mutation rate evolution for the rate of adaptation within the population. Finally, we describe the approximations we have made throughout our analysis and the resulting limitations of our approach.

### Approach to the stable mutation rate

Consider a situation in which the population starts with a mutation rate *U*_0_ ≠ *Û* and fixes a sequence of modifier alleles with mutation rates *U*_1_, *U*_2_,…, *U_n_*. What are the typical values of this mutation rate trajectory, under the hypothesis that each step was favored by selection?

In both the successional mutations and clonal interference regimes, we found that *Û* depends only on *N*, *S_b_*, and ∈. This implies that the mutation rate will always evolve towards this unique stable point: *U_n_* → *Û* However, the manner in which the population approaches *Û* can be strongly influenced by clonal interference.

In the successional mutations regime, selection favors a monotonic approach to *Û* Consider, e.g., a mutation rate *U_0_* < *Û* In this case, we have shown that mutators will be able to fix provided that the new mutation rate *U_m_* = *rU* lies in the range *U_0_* < *U_m_* < *Û* When one of these alleles fixes, the new mutation rate will be given by *U_1_* = *U_m_*, and the process will repeat itself. Mutators will still be favored to fix provided that *U_1_* < *U_m_* < *Û* which will lead to a *U_2_* > *U_1_*, and so on. Thus, evolution will favor a monotonic sequence of mutator alleles until *U_n_* = *Û* after which the mutation rate will be stable to further changes. An exactly analogous conclusion holds if *U_0_* starts above *Û*: here selection will tend to fix antimutator alleles that lie in the range *Û* < *U_m_* < *U_i_* until *U_n_* = *Û* In both cases, mutator or antimutator alleles that “overshoot” the stable mutation rate are always disfavored. This is true even if such alleles would move the mutation rate closer to the stable rate; e.g., an allele that takes a population from 0:1*Û* → 1:1*Û* would not be positively selected.

**FIG. 4.**
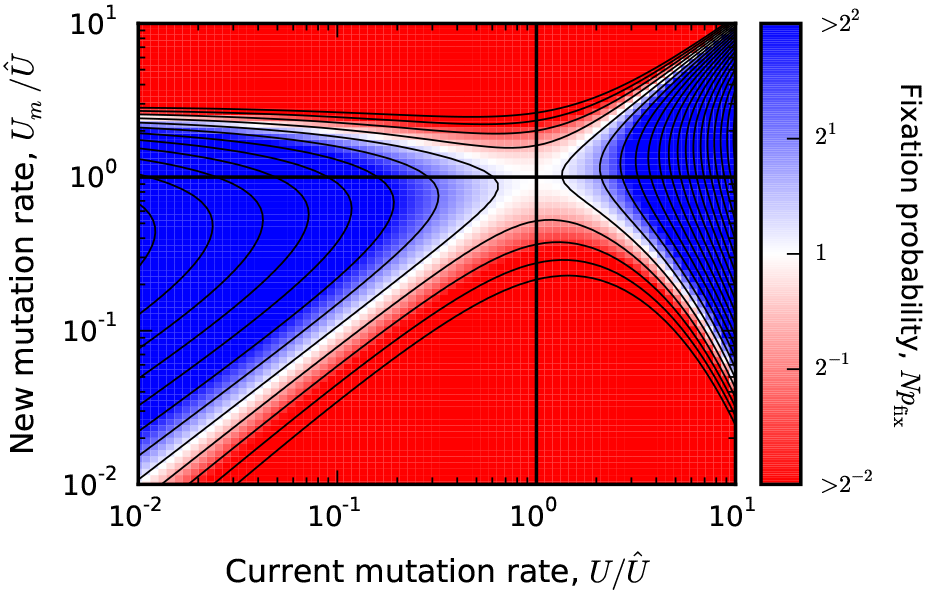
The predicted fixation probability of a modifer allele when *Ns_b_* = 10^6^ and ∈ = 10^-6^. Grid points are colored according to the value of *N*_*p*fix_ from Eq. (38), and are capped at a maximum value of |log_2_ *N*_*p*fix_| = 2 to maintain contrast.For comparison, the solid lines denote twenty log-spaced contours that range from 10^-1^ to 10^4^.

When clonal interference is present, the approach to the stable mutation rate is more complex. This is easiest to see if we rewrite Eq. (20) in terms of *U_m_*, *U*, and *Û*. Provided that 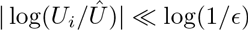, the quantities *τ*_est_(*U*) and *q*(*U*) will stay roughly constant over the relevant range of mutation rates, and we can approximate 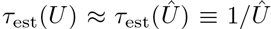. This yields a simple heuristic formula,

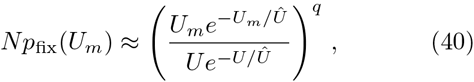

which depends only on the scaled mutation rates *U*/*Û*, *U_m_*/*Û*, and *q*. A more accurate (but more cumbersome)version can be obtained by substituting the full expressions for *τ*_est_(*U*) and *q*(*U*) into Eq. (38). We illustrate this full expression in Fig. 4 for a particular combination of *N s_b_* and ∈, although the important qualitative features are already contained in Eq. (40).

If the population starts at a mutation rate *U*_0_ ~ Û,then only modest changes in the mutation rate will be favorable.However, if the population starts at a mutation rate *U*_0_ ≪ *Û*, Eq. (40) shows that mutators will be positively selected provided that 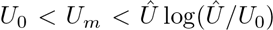.While the most strongly beneficial modifier has *U_m_* = *Û*, the range of favored mutators also includes values of *U_m_* that are *larger* than *Û* by a factor log(*Û*/*U*_0_) ≫ 1. This means that selection can favor mutator alleles that overshoot the stable mutation rate by a substantial amount.If a mutator allele of this maximal strength fixes, the new mutation rate will be 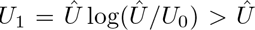, and selection will immediately favor the fixation of antimutator alleles, despite the fact that the location of *Û* has not changed (see Fig. 5). If *U* ≫ *Û*,these antimutator alleles can overshoot the stable mutation rate in the other direction,by a factor 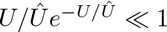. In the example above,this could then lead to *U*_2_ = *U*_0_log(*Û*/*U*_0_), which is larger than the initial mutation rate *U*_0_ but still much smaller than *Û*.Thus while the population will still approach the stable mutation rate over time, this approach does not have to be monotonic, even in the absence of epistasis or environmentalvariation. Moreover, the functional form of Eq. (40) implies that this behavior is highly asymmetric:an antimutator allele can overshoot *Û* by a larger amount (and with a larger *N*_*p*fix_)than a mutator allele with the same value of | log(*U*/*Û*)j. This can also be seen in Figs 4 and 5.

**FIG. 5.**
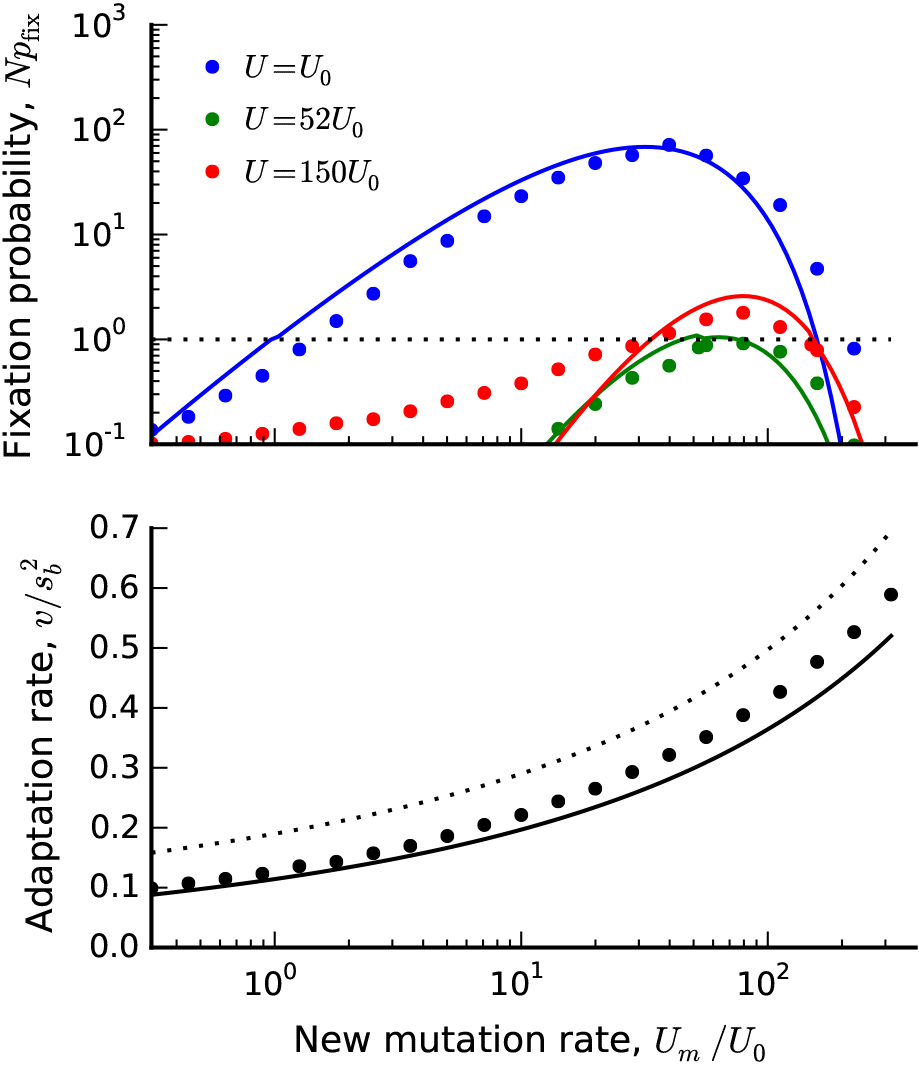
(Top) A vertical “slice” of Fig. 4 for three different values of *U*. Symbols denote the results of forward time simulations with *NU*_0_ = 5780, and *s_d_* = 5*s_b_*; other parameters are the same as Fig. 4. Solid lines denote theoretical predictions from Eq. (38). (Bottom) The scaled rate of adaptation as a function of the mutation rate for the same set of parameters.Symbols denote the results of forward-time simulations, the solid lines show the theoretical predictions obtained by solving Eqs. (31) and (32) numerically, and the dashed line shows the asymptotic formula, 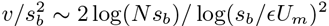.

### Application to a long-term evolution experiment in E. coli

Several studies have observed the spread of mutator alleles in microbial populations in the laboratory. One of the best-studied examples is Lenski’s long-term evolution experiment (the LTEE), where 12 replicate populations of *E. coli* have been propagated constant conditions for more than 60,000 generations (Lenski et al., 1991, 2015). Mutator phenotypes have fixed in 6 of the 12 populations over the course of this experiment, and have typically increased the mutation rate by ~ 100-fold. The rate of adaptation in the mutator populations increased less than 2-fold, suggesting that clonal interference plays an important role (Wiser et al., 2013). In one of the 12 replicates, the dynamics of mutation rate evolution have been studied in more detail. Wielgoss et al. (2013) have shown that, soon after the fixation of the mutator allele, an antimutator phenotype fixed, which lowered the mutation rate by a factor of two. The antimutator phenotype had two independent origins, suggesting that both it and the original mutator allele were favored by selection. Given these findings, it is interesting to compare various explanations for this behavior in the context of our theoretical analysis above.

One potential explanation for these findings is suggested by the “overshooting” behavior illustrated in Figs 4 and 5. Although the precise evolutionary parameters of these populations are not known, Fig. 5 shows that it is at least feasible to overshoot the optimum by a factor of 2 when the population starts at a mutation rate of *U*_0_ ~ 0.01*Û*. Note, however, that *N*_*p*flx_ is not necessarily large in the reverse direction, so it remains to be seen whether this effect could efficiently select for lower mutation rates on the rapid timescales observed.

Another potential explanation for the mutation-rate reversal, initially suggested by Wielgoss et al. (2013), is that long-term epistasis reduced the effective advantage of hitchhiking later in the experiment, thereby shifting the location of *Û* below its initial value. In the context of our model, this epistasis can manifest itself in one of two ways. First, it could reduce ∈, reflecting a diminished supply of beneficial mutations (or an increasing supply of deleterious mutations) as the population adapts. Our formula in Eq. (25) suggests that rather drastic reductions in ∈ are required to cause a 2-fold reduction in *Û*. Alternatively, diminishing-returns epistasis could reduce the magnitude of *s_b_*, while leaving the beneficial mutation rate unchanged. Equation (25) suggests that this has much stronger effect on *Û* Wiser et al. (2013) have recently proposed and fit a concrete model for how *S_b_* declines with fitness in the LTEE. These estimates suggest that *s_b_* declines by ≲20% in the 4,000 generations that separate the mutator and antimutator alleles. This would seem to be too small to account for a≳50% reduction in *Û*, although the magnitudes are sufficiently close that a careful comparison with simulations is required. This would be an interesting avenue for future work.

A third potential explanation is a direct fitness benefit to either the mutator or antimutator allele. While such benefits are difficult to measure directly (due to the confounding effects of the deleterious load), they can be incorporated into our model in a straightforward way. We can simply replace *U_d_*(*r* − 1) → *U_d_*(*r* − 1) − *s_m_* in our formulae for *p*_flx_(*r*), where *s_m_* is the direct benefit of the modifier allele. Since the effect of the deleterious load is typically of order *Û* < *s_b_*, even a small direct benefit (*s_m_* < *s_b_*) can shift the balance between hitchhiking and load in important ways.

### The evolution of mutation rates and the average rate of adaptation

Because we have assumed that deleterious mutations are purgeable, increasing *U* always increases the rate of adaptation, even though these increases may be small in the presence of clonal interference. Our results therefore suggest that mutation rates will evolve toward a stable mutation rate that is less than what would be optimal for the population (which, in this case, is technically *U*_opt_ = ∞). Of course, as we increase mutation rates, the assumption that deleterious mutations are purgeable will eventually fail. Somewhere above this point, there will be an optimal mutation rate *U*_opt_ < ∞ that maximizes the rate of adaptation (see, e.g. Orr (2000)). To confirm that the stable mutation rate is indeed below *U*_opt_, we must verify that *Û* is still in the purgeable deleterious mutations regime. For the example in Fig. 5, we can verify this directly, since the rate of adaptation continues to increase as the mutation rate passes through *Û*. This behavior will hold more generally provided that the typical cost of a deleterious mutation, *s_d_*, is much larger than *Û*. Since *Û* is typically smaller than *S_b_* in the regimes that we consider, this will indeed be the case whenever *s_d_* ≳ *s_b_*.

In these cases, the stable mutation rate is below the optimal mutation rate, which implies that the dynamics of the evolutionary process can sometimes favor changes in mutation rate that slow the adaptation of the population. Antimutator alleles can be favored even when their fixation will ultimately reduce the overall rate of adaptation. Conversely, mutator alleles can be disfavored even when their fixation would have increased the rate of adaptation.

Although our results are limited to the purgeable regime, a similar distinction between *Û* and *U*_opt_ may also apply even when deleterious mutations are no longer purgeable. We cannot prove this conjecture in our present framework, but it is an interesting hypothesis for future work.

### Limitations to our analysis

We have made a number of key assumptions throughout our analysis. Most crucially, we have assumed that deleterious mutations are purgeable. For this to be true, two conditions must be met: the deleterious mutations cannot fix, nor can they affect the overall rate of adaptation by reducing the fixation probabilities of beneficial mutations. This is a slightly stronger condition than the “ruby in the rough” approximation used by earlier authors (Charlesworth, 1994; Peck, 1994). Our previous work on the effects of deleterious mutations in adapting populations provides a detailed analysis of when these conditions will hold, and of the effects of deleterious mutations on adaptation when the purgeable assumption fails (Good and Desai, 2014). This earlier work suggests that deleterious mutations are likely to be purgeable in most microbial evolution experiments, though recent experimental work hints that this may not always be the case (McDonald et al., 2016). Here we first summarize the conditions under which deleterious mutations are purgeable, and then describe the potential effects of deleterious mutations when this condition is violated.

In our earlier work, we showed that deleterious mutations with fitness cost *s_d_* will typically not fix provided that *s_d_* ≫ 1/*T_c_*, where *T_c_* ~ 1/*qτ*_est_ is the coalescence timescale (Good and Desai, 2014). Since *s_b_* ≫ 1/*T_c_* in all the regimes that we consider, this will be true provided that *s_d_* is not much smaller than *s_b_*. Deleterious mutations can also hinder the fixation of beneficial mutations that arise in less-fit backgrounds. However, provided that s*_d_* ≫ *U_d_*, deleterious variants will be rapidly eliminated from the population, and most genetic backgrounds will be free of deleterious mutations. Assuming typical values of *U_d_* ~ 10^-4^ and *s_d_* ~ 10^-2^ − 10^-1^ for microbial evolution experiments (Wloch et al., 2001), this condition will often be met.

When deleterious mutations have small enough fitness costs that they are not purgeable, our quantitative results all break down. However, sufficiently weakly deleterious mutations cannot affect mutator or antimutator dynamics on the relevant timescales, so they are effectively neutral from the point of view of mutation rate evolution. On the other hand, sufficiently strongly deleterious mutations are purgeable. Thus it is only some range of deleterious mutations between these limits whose effects are both important to mutation rate evolution and not described by our analysis. Since these mutations are less deleterious than purgeable mutations, they must by definition be less unfavorable to mutator alleles (and less favorable to antimutators). Thus they do not affect the fate of a modifier of mutation rates as strongly as a corresponding purgeable mutation would. This suggests that we can qualitatively understand the effects of non-purgeable deleterious mutations in adapting populations by weighting them less heavily than purgeable ones, so that the total effective deleterious mutation rate is actually somewhat less than *U_d_*. Of course, this is only a rough intuition. To analyze the effects of non-purgeable deleterious mutations more fully, we need to include them in our traveling wave framework. Our earlier work explains how these mutations affect *f*(*x*), *w*(*x*), *x_c_*, and *v*, which provides a potential starting point for these calculations (Good and Desai, 2014). More generally, we may wish to consider the evolution of mutation rates in the limit where non-purgeable mutations become so important that the population no longer adapts, and instead Muller’s ratchet plays a crucial role. These are interesting but complex directions for future work.

In addition to these key limitations of our analysis, we have also made a number of more technical assumptions. For example, we have assumed that beneficial mutations all provide the same fitness benefit *s_b_*. In reality, beneficial mutations will have a range of different fitness effects, drawn from some distribution of fitness effects. However, earlier work has shown that in this case the evolutionary dynamics can be summarized using a single effective beneficial fitness effect and corresponding effective beneficial mutation rate (Desai and Fisher, 2007; Fisher, 2013; Good et al., 2012). Thus our analysis can be applied to this situation using the appropriate effective *s_b_* and *U_b_*.

In our analysis of clonal interference, we focused on a regime in which 1 ≪ log(*s_b_*/*U_b_*) ≪ 2log(*N s_b_*) ≪ log^2^(*S_b_*/*U_b_*). The latter condition implies that clonal interference is not exceptionally strong, and is often a good approximation for microbial evolution experiments at wildtype mutation rates. For some large effect mutators, we start to approach the boundary of this regime, and our quantitative expressions become less accurate (see Fig. 5). In principle, we could extend our analysis to the “high-speed” [2log(*Ns_b_*) ≫ log^2^(*s_b_*/*U_b_*)] or “mutational-diffusion” regimes [*U_b_* ≫ *s_b_*] by using analogous solutions for *f*(*x*) and *w*(*x*) derived by Fisher (2013) or Hallatschek (2011). We leave this for future work.

Throughout our analysis, we have assumed that modifier effect sizes can be large but not exceptionally so [e.g., in the clonal interference regime, |log(*r*)| ≪ log(*s_b_*/*U_b_*)]. Since *U_b_* ≪ *s_b_*, this includes many realistically large mutators of order *r* ~ 100, and in practice, our expressions appear to work reasonably well even when log(*r*) ~ log(*s_b_*/*U_b_*) (see Fig. 3). For sufficiently large *r*, we may also encounter situations where modifiers switch from one regime to another (e.g., from the clonal interference to successive mutations regime, or from *U_b_* ≪ *s_b_* to *U_b_* > *s_b_*). Our quantitive predictions for *N*_*P*flx_ break down in all of these cases. However, the evolutionarily stable mutation rates are defined by the behavior of *N*_*p*flx_ in the local neighborhood of *r* ≈ 1, so the location of *Û* is not influenced by this problem. If the population starts sufficiently far from *Û* then the fixation probabilities of the first few *U*_i_ are not well-described by our analysis, but we know that the mutation rate must still approach *Û* (in a possibly non-monotonic manner). Eventually, the mutation rate will become close enough to *Û* that our expressions start to apply, and the remaining steps of the mutation trajectory can be predicted.

Finally, we have assumed throughout that modifier mutations are sufficiently rare that their fates are determined independently. In other words, we have neglected clonal interference *between* different modifier mutations. For deleterious or weakly beneficial modifiers, this will often be a good approximation. However, for strongly beneficial modifiers, this requires that the establishment time 1/*N* _*μp*_flx__(*r*) between successive mutators is large compared to the fixation time. In the clonal interference regime, *T*_fix_ ~ log(*s_b_*/*U_b_*)/*s_b_*, and this condition becomes *μ* ≪ *S_b_*/log(*s_b_U_b_*)*N*_*p*flx_(*r*). This can sometimes be violated for strongly beneficial mutator alleles with a large target size (e.g., loss-of-function mutations in multiple genes). In these cases, our quantitative predictions become inaccurate, although the overall direction of selection [i.e., whether *N*_*p*flx_ > 1 or *N*_*p*flx_ < 1] will remain unchanged.

## Conclusions

Our analysis has explained how the interplay between beneficial and deleterious mutations in adapting populations creates indirect selection pressures on modifiers of mutation rates, and we have shown how these indirect selection pressures affect the fates of mutator and antimutator alleles. Our analysis of the successional mutations regime follows the logic of earlier work (Andre and Godelle, 2006; Desai and Fisher, 2011), balancing the probability a new beneficial mutation arises in a mutator or antimutator background with the effects of the modifier allele on the accumulation of deleterious load. We have also studied rapidly adapting populations where clonal interference is widespread. Our approach to this question builds on the traveling wave framework we and others have recently introduced (Fisher, 2013; Good and Desai, 2014; Good et al., 2012; Hallatschek, 2011; Neher and Shraiman, 2011; Neher et al., 2010). In this framework, analyzing the fate of an allele modifying mutation rate is similar in spirit to calculating the fixation probability of any lineage: we must solve the same equation for *w*(*x*), except instead of a mutation changing the fitness *x*, it changes the mutation rate *U*. This general framework can also be applied to analyze indirect selection pressures that act on other modifiers of the evolutionary process. For example, we could analyze the fate of a mutation that changes the distribution of fitness effects of new mutations by solving for *w*(*x*) for an allele that modifies *ρ*(*S*). Our analysis does not need to be limited to adapting populations, since the traveling wave framework applies whenever interference selection is widespread, even if the population is not adapting on average. Of course, in practice, some modifiers and parameter regimes will lead to equations for *w*(*x*) that are analytically tractable, while others will not. Thus further work is needed to more fully understand both the limits and promise of this approach.

## ACKNOWLEDGMENTS

We thank Ivana Cvijović for useful discussions and helpful comments on the manuscript. Simulations in this article were run on the Odyssey cluster supported by the FAS Division of Science Research Computing Group at Harvard University. This work was supported in part by the Simons Foundation (grant 376196), grant PHY 1313638 from the NSF, and grant GM104239 from the NIH.

## Appendix A: Exact solution of Eq. (3)

In this section, we evaluate the integral

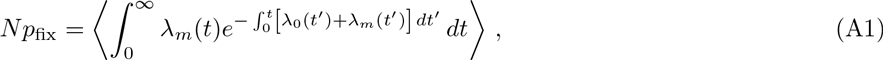

where 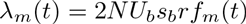 and 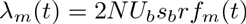, and *f*_m_(*t*) satisfies the Langevin dynamics

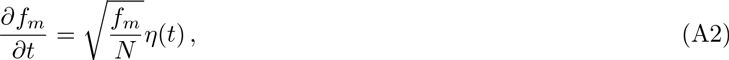

with *f*_*m*_(0) = 1/N. Using integration by parts, we can rewrite this integral as

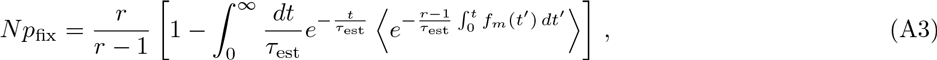

where we have used the short-hand 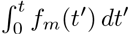. To evaluate this integral, we must first solve for the generating function H(*y, t*) for the weight 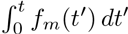, which is defined by

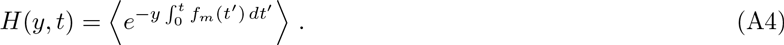

This can be done using techniques described in Weissman et al. (2009). Briefly, we first introduce the joint generating function,

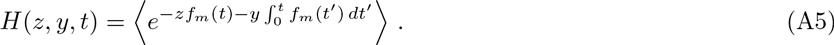

By Taylor expanding *H(z, y, t + dt)* and using the Langevin dynamics in Eq. (A2), we can show that *H(z, y, t)* obeys the partial differential equation

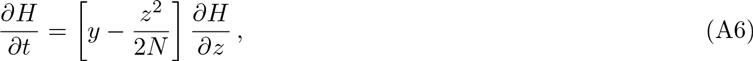

which can be solved using the method of characteristics. The characteristic curves satisfy

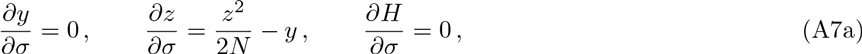

and the solution is given by

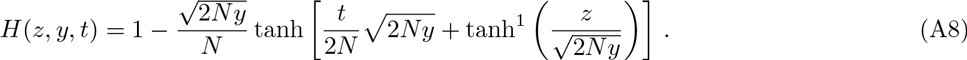

The marginal generating function for 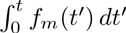 can then be obtained by setting *z* = 0:

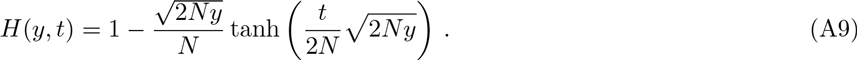

Subsituting this expression into the integral in Eq. (A3), we find that

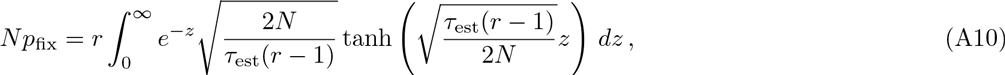

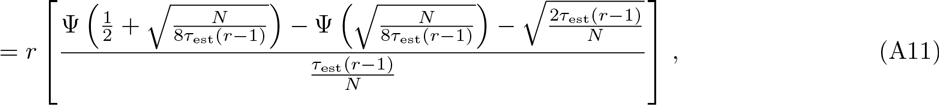

where ψ(*z*) is the digamma function. Asymptotic expansions for small and large arguments yield

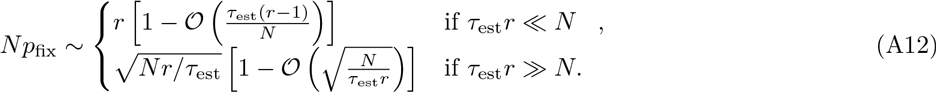

## Appendix B: Derivation of Eq. (38)

To obtain an expression for *N_Pfix_(r)*, we must evaluate the integral

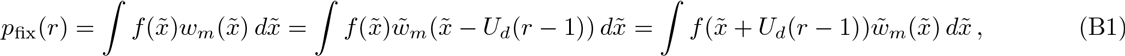

using the expressions for 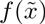 and 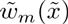 in Eqs. (29) and (34) in the main text. This is straightforward, although the algebra is somewhat tedious. Since 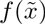 and 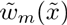 are piecewise functions, the form of the integrand will depend on whether *x_c_ − U_d_(r − 1)* is greater or less than *x_cm_* and *x_cm_ − S_b_*.

We first consider the case where *x_c_ − U_d_(r − 1) > x_cm_*. In this case, the mutator interference threshold is reached before the nose, so that

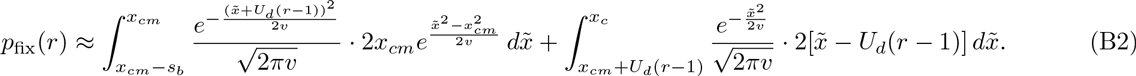

After multiplying both sides by 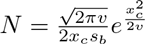, we obtain

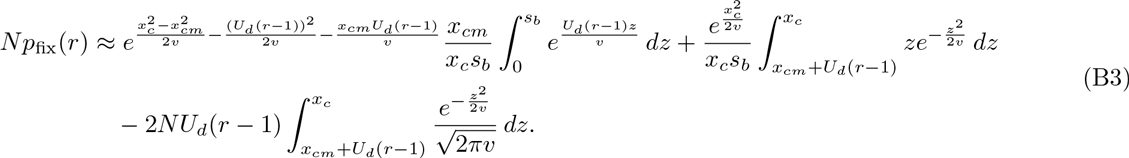

Evaluating the remaining integrals and substituting 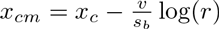, we find that

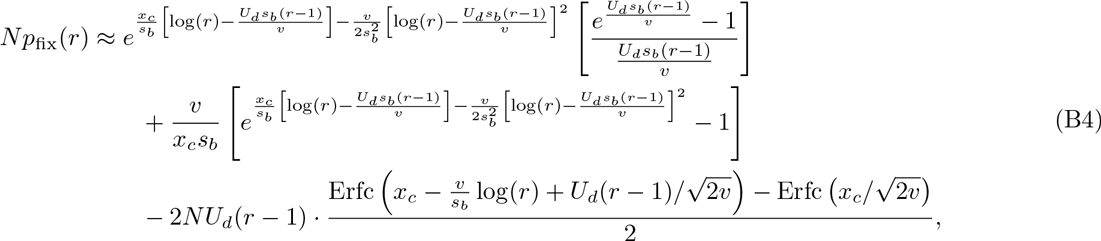

where 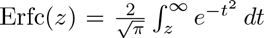. The last two terms are only relevant for extremely high deleterious mutation rates where *U_d_(1 − r) ∼ x_c_*. After neglecting these terms, we obtain Eq. (11) in the main text.

Now we consider the case where *x_c_ − U_d_(r − 1) < x_cm_*. In this case, the edge of the fitness distribution is reached before the mutator interference threshold. If *x_c_ − U_d_(r − 1) < x_cm_ − S_b_*, then the edge of the fitness distribution never even makes it to the region where 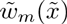 is positive, and *p_fix_(r) ≈ 0*. On the other hand, if *x_c_ − U_d_(r − 1)* lies between *x_cm_* and *x_cm_ − S_b_*, then

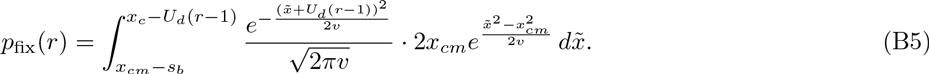

Multiplying by 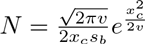 and evaluating the integrals, we obtain

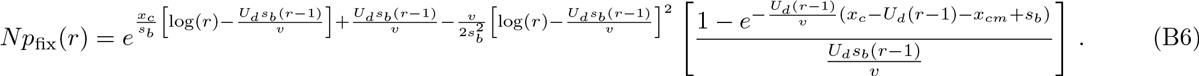

In practice, the differences between Eqs. (B4) and (B6) do not become very pronounced until *N_Pfix_(r)* is already quite small. For simplicity, we will therefore use Eq. (B4) for the full range of *r*.

## Appendix C: Forward-time simulations

We validate several of our key approximations by comparing our predictions with the results of forward-time simulations similar to the standard Wright-Fisher model. In these simulations, the population is divided into fitness classes depending on the number of beneficial mutations (*n_b_*) and the number of deleterious mutations (*n_d_*) that each individual possesses. The simulation starts with a homogeneous population of *N* individuals with *n_b_* = 0 and *n_d_* = 0. To form the next generation, each individual produces a Poisson number of identical offspring with mean *C_t_(1 + s_b_n_b_ − s_d_n_d_)(1 − U_d_ − U_b_)*, a Poisson number of beneficial offspring (*n_b_ → n_b_ + 1*) with mean *C_t_(1 + s_b_n_b_ − s_d_n_d_)U_b_*, and a Poisson number of deleterious offspring (*n_d_ → n_d_ + 1*) with mean *C_t_(1 + s_b_n_b_ − s_d_n_d_)U_d_*, where *C_t_* is a constant recalculated at each generation to ensure that the expected number of offspring for the entire population is *N*. Starting from the initial population, all simulations are allowed to “burn-in” for *Δt* = 5 x 10^4^ generations before any measurements are made.

To measure the fixation probability of a modifier allele, we modify this basic algorithm so that new modifier offspring are produced from wildtype individuals at rate *μ*. These modifier lineages reproduce the same way as wildtype individuals except with *U_b_ ₒ rU_b_* and *U_d_ ₒ rU_d_*. Further modifications to the mutation rate or reversion to the wildtype are not allowed. To minimize the effects of initial conditions, *μ* is artifically fixed at zero until the burn-in period has elapsed. We then record the number of generations *T* between the end of the burn-in period and the fixation of the modifier phenotype. In the limit that *μ* ₒ 0, this is related to the fixation probability through the relation

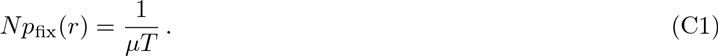

To ensure that *μ* is chosen to be small enough for Eq. (C1) to apply, we repeat this process *M* = 60 times with a sequence of modifier mutation rates *μ*_1_, *μ*_2_,…, *μ_M_*, and a sequence of fixation times *T*_1_, *T*_2_,…, *T_M_*. The mutation rate at step *i* is chosen based on the previous *T*_*i*−1_, so that the predicted value of *T_i_* is 10^4^ generations:

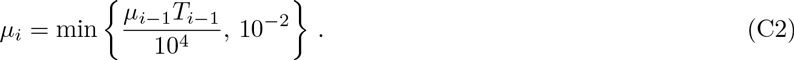

The value 10^4^ is chosen because, for the parameters considered here, 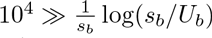, and the mutation rates are capped at 10^−2^ to minimize the correlated effects of Muller’s ratchet (Neher and Shraiman, 2012). The sequence is started with *μ*_1_ = 10^−4^, and is allowed to “burn-in” for 10 iterations before *T_i_*’s are recorded. The fixation probability is calculated from the maximum likelihood estimator,

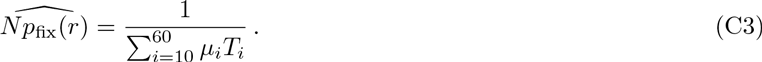

A copy of our implementation in Python is available at https://github.com/benjaminhgood/mutator_simulations.

